# Integrating single-cell and bulk transcriptomes identifies neuronal heat-stress-associated candidate genes in a reef-building coral

**DOI:** 10.64898/2026.10.05.756843

**Authors:** Ruiqi Li, Martin Tresguerres, Carly Kenkel

## Abstract

Cell-type-specific molecular responses are obscured in bulk RNA sequencing, whereas generating matched single-cell datasets across environmental conditions remains costly and technically challenging, particularly in non-model organisms. Here, we present a workflow that integrates existing bulk and single-cell RNA-seq resources to prioritize cell-type-associated environmental-response genes. We demonstrate the workflow by examining heat-stress-associated neuronal genes in the reef-building coral *Stylophora pistillata*. Neuronal genes were first identified based on cell type-specific expression in single-cell transcriptomic data and then intersected with differentially expressed genes from bulk RNA-seq datasets under heat-stress. In parallel, we applied a deconvolution approach to infer cell type-specific expression changes directly from bulk data, independently capturing neuronal transcriptional responses. This integrated pipeline reduced thousands of heat-responsive genes previously identified in bulk RNA-seq studies to nine heat-associated, neuron-enriched candidates. Eight of the nine candidates showed their highest mean expression in the same neuronal subcluster. Targeted sequence-based characterization supported the identification of a TRPA-family ion channel, two calmodulin/centrin-family Ca²-binding proteins, and a Class A/rhodopsin-like GPCR of unresolved ligand specificity. Two additional candidates were predicted secreted proteins belonging to conserved coral protein families, while the molecular functions of the remaining candidates remained unresolved. Their coordinated regulation suggests heat-associated remodeling of neuronal sensory and intracellular signaling processes. Our workflow provides a scalable framework for extracting cell type-associated molecular responses from existing transcriptomic resources and prioritizing candidates for further investigation. More broadly, it offers a generalizable strategy for investigating cell type-specific gene expression responses across organisms, cell types, and environmental conditions without requiring new single-cell datasets.

## Introduction

Animals must detect and process internal and external cues to coordinate adaptive physiological and behavioral responses to environmental variation. The underlying mechanisms are inherently heterogeneous and are mediated by distinct cell populations, each relying on a specific transcriptional program^1^. This cellular heterogeneity critically influences tissue, organ, and organismal function, yet is obscured in bulk RNA-sequencing (bRNAseq)^2^. In contrast, single-cell RNA sequencing (scRNAseq) resolves cell type-specific transcriptional programs and their contributions to organismal responses^3^. However, generating single-cell datasets remains costly and technically challenging, particularly for environmental studies involving multiple experimental conditions and extensive replication^4^. Consequently, bRNAseq remains widely used for studying transcriptional responses across diverse species and environmental conditions^5^.

Bulk transcriptomes average signals across cell populations and can mask distinct, or even opposing, cell-type-specific responses to environmental stress. This challenge is especially relevant in reef-building corals, which lack organs and instead have diverse specialized cell types intermingled within the same tissues. Unlike animals in which organs such as the brain, gills, or liver can be dissected to enrich for particular cell types and associated physiological functions, whole-coral samples combine transcriptional signals from many functionally distinct cell populations. For example, responses to thermal stress may emerge from the coordinated activity of multiple specialized cell types, such as neurons that mediate environmental sensing, endodermal cells that regulate algal symbiont dynamics, and calicoblasts that control skeletal growth. Resolving transcriptional changes within these individual cell types would enable a more mechanistic understanding of how corals detect and respond to environmental stressors that increasingly threaten coral populations globally^6^.

In humans and other model systems, computational deconvolution approaches are increasingly used to integrate single-cell reference data with bRNAseq and infer cell-type-associated signals from heterogeneous tissues^7,8^. The recent development of single-cell transcriptomics for reef-building corals^9^, together with the extensive body of environmental bRNAseq studies^10^, provides an opportunity to integrate these independently generated datasets and resolve cell-type-associated responses to environmental change. Such integration could provide cellular context for previously characterized transcriptional responses without requiring new single-cell experiments for every environmental condition. Beyond resolving responses at cell-type level, cellular context can also aid the functional interpretation of transcriptomic data in non-model organisms, which is hindered by the limitations of homology-based annotation^11^. Homology-based approaches often fail to capture lineage-specific genes and can propagate uncertain functional assignments from distantly related organisms. Cellular context provides an additional layer of biological information that can help prioritize candidate genes for downstream functional investigations, despite uncertain or uncomplete annotation.

We address those challenges by integrating bRNAseq and scRNA datasets from the reef-building coral *Stylophora pistillata,* to illustrate a workflow for identifying cell-type-associated responses to environmental stress and prioritizing candidate genes related to targeted cell types and functions in non-model organisms. Importantly, *S. pistillata* also has a highly complete genome assembly and comparatively well-developed genomic resources, providing a strong foundation for integrating single-cell and bulk transcriptomic datasets. We illustrate this approach (Fig. 1) for neuronal responses to thermal stress, combining single-cell-derived neuron-specific gene signatures with bRNAseq data from two independent thermal stress exposure studies. We also used Bayesian deconvolution as a complementary approach to identify heat-responsive genes associated with neuronal cell populations. Concordance across these complementary analyses was then used to prioritize candidate neuronal genes associated with heat stress. Candidate proteins were subsequently characterized using complementary sequence-based analyses to evaluate and refine their functional annotations. This framework can, in principle, be applied to other cell types and environmental conditions represented by corresponding single-cell and bulk transcriptomic resources.

**Figure 1.**
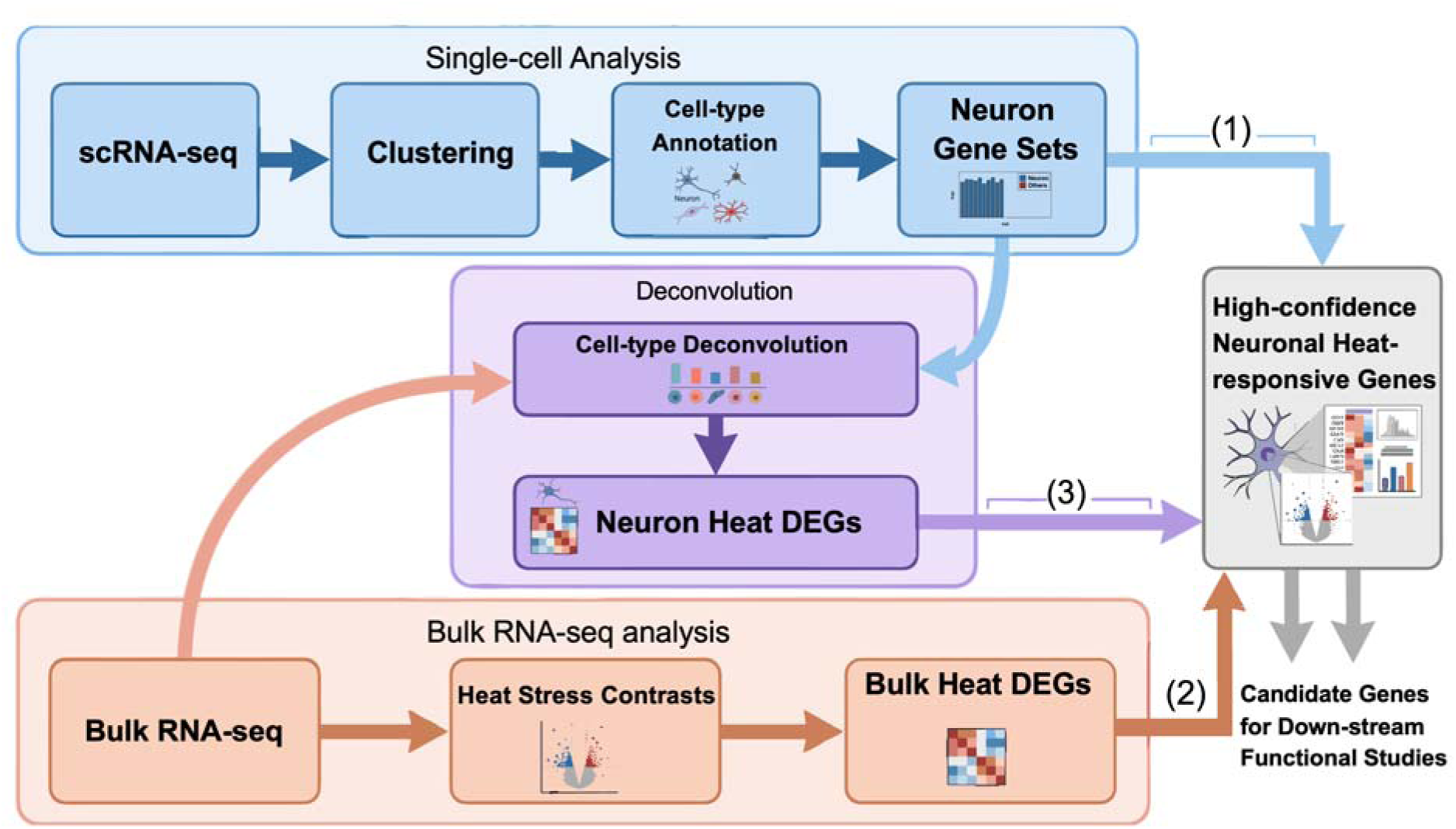
Overview of analytical workflow for identification of neuronal heat responsive genes. Single cell and bulk RNA sequencing analyses were integrated to identify high confidence neuronal heat responsive genes: 1) Identification of neuronal genes 2) DEG analyses on heat stressed corals 3) Deconvolution. Genes supported by both approaches and both studies were retained as consensus candidates.

## Methods

### 1. Genome Reference and Gene Model Extension

The chromosome-level genome assembly of *Stylophora pistillata* jaStyPist1.1 (GCA_964205215.1) was used as the host reference. The predicted annotation comprises 30,756 gene models^12^. Genome completeness was assessed using BUSCO with the Cnidaria odb12 lineage dataset and was 98.6%^13^. Because 3’ UTRs are frequently incompletely annotated in non-model species, gene models were extended using 10X Genomics Chromium scRNAseq data following Levy et al.^9^ (2021). Specifically, reads were aligned to the *S. pistillata* genome using STAR^14^. The resulting BAM files were used to define genomic intervals enriched for 10X Genomics reads using MACS2^15^. Stranded genomic intervals were used to extend the annotated 3’ ends of gene models. The extended gene models were used for subsequent scRNAseq and bRNAseq analyses. To enable competitive mapping of potential symbiont reads, the genome of the dinoflagellate algal symbiont *Symbiodinium microadriaticum* pySymMicr2.1 (GCA_977109205.1) was concatenated with the *S. pistillata* genome. The combined host-symbiont reference was indexed using Cell Ranger 10.0.0^16^ and HISAT2 2.2.1^17^ for downstream scRNAseq and bRNAseq mapping, respectively.

### 2. Single-Cell RNA-Seq Data Processing and Clustering

scRNA-seq data generated using 10X Genomics Chromium technology from adult *S. pistillata* were retrieved from PRJNA1223412^18^. Analyses were performed on the University of Southern California Center for Advanced Research Computing (CARC) high-performance computing cluster. Raw data were processed using Cell Ranger 10.0.0^16^ with BAM file generation enabled. The filtered gene-barcode matrix was imported into R 4.4.2^19^ and analyzed using Seurat 5.3.0^20^. Quality control filtering was applied to remove low-quality cells and potential doublets. Cells were retained if the number of detected genes (nFeature_RNA) was between 400 and 3,500 and the total UMI count (nCount_RNA) was below 15,000. The percentage of mitochondrial transcripts was not used as a filtering criterion because the reference annotation did not include mitochondrial chromosome annotations. Filtered data were normalized using Seurat’s NormalizeData function with the LogNormalize method and a scale factor of 10,000. Highly variable features were identified using FindVariableFeatures with the variance stabilizing transformation method, selecting the top 3,000 genes. Data were scaled and centered using ScaleData with default parameters. Principal component analysis was performed on the scaled expression matrix of variable genes using RunPCA, computing 50 principal components. Based on an elbow plot, the first 20 principal components were retained for downstream analyses. A shared nearest neighbor graph was constructed using FindNeighbors with the first 20 principal components. Graph-based clustering was performed using the Louvain algorithm implemented in FindClusters with a resolution of 0.2. Two-dimensional visualization was generated using UMAP with RunUMAP using the first 20 principal components.

### 3. Cell Cluster Annotation

Cluster marker genes were identified using the FindAllMarkers function in Seurat 5.3.0. Marker genes were identified using the Wilcoxon rank-sum test, with a minimum fraction of cells expressing the gene in either group (min.pct = 0.25), and a minimum log fold-change threshold (logfc.threshold = 0.25). Marker genes were annotated with gene names and functional descriptions using eggNOG-mapper^21^. Because most top-ranked marker genes were unannotated, top marker gene identity alone was insufficient for reliable cell-type assignment. To address this limitation, a top-down orthology-based strategy was applied using previously published cell-type annotations from *S. pistillata* (Levy et al., 2021)^9^ generated with an earlier scaffold-level genome assembly. For each cell type reported in Levy et al. (2021)^9^, the top marker genes were retrieved. OrthoFinder^22^ was used to identify orthologous relationships between genes from the previous genome version and the chromosome-level genome assembly used in this study. Expression of mapped orthologs was visualized across clusters using UMAP feature plots and dot plots in Seurat. Based on the expression patterns of these orthologous markers, eight major cell types were annotated: neuron, germline, immune, gastrodermis, epidermis, alga-hosting cells, calicoblast, and gland. Clusters lacking strong or specific marker expression were assigned as “other.” As a complementary approach, gene names and keywords derived from marker genes reported in Levy et al. (2021) were used to identify corresponding genes in the updated gene models of the chromosome-level genome assembly. For example, neuronal markers such as GABBR2, NPTX1, and genes annotated with the keyword “Synapsin” were used to retrieve candidate genes in the annotation. Expression of these candidate genes was visualized across clusters using UMAP feature plots and dot plots. These results further supported the final cluster annotations.

### 4. Identification of Neuronal Genes

To identify genes associated with neuronal expression from the annotated single-cell dataset, two methods were applied: differential expression analysis to identify genes highly expressed in neurons relative to other cell types (*neuron_highexpr*), and a proportional expression metric to identify genes for which most detected transcripts occurred in neurons (*neuron_predominant*). First, differential expression analysis was performed using the FindMarkers function in Seurat 5.3.0 to compare neuronal cells against all other cells. Genes were classified as highly expressed in neurons if they met the following criteria: logfc ≥ 1.0 and adjusted p-value < 0.05 using Bonferroni correction. This gene set is hereafter referred to as *neuron_highexpr*. Second, to identify genes predominantly expressed in neurons, a neuron expression fraction was calculated for each gene. Total UMI counts were summed across all neuronal cells (neuron_total_counts) and across all cells in the dataset (all_total_counts). The neuron expression fraction was defined as neuron_expr_fraction = neuron_total_counts / all_total_counts. Genes with neuron_expr_fraction ≥ 0.80 were classified as predominantly expressed in neurons. This gene set is hereafter referred to as *neuron_predominant*.

### 5. Bulk RNA-Seq Data Processing and Differential Expression Analysis

Two previously published bulk RNA-seq datasets were analyzed. The Voolstra et al. (2021)^23^ dataset comprised 83 samples representing four coral populations exposed to three temperature treatments (30°C control, 33°C intermediate, and 36°C heat stress). The ICN population represents the most heat-sensitive population in this dataset. The Savary et al. (2021)^24^ dataset included 80 samples from short-term (S) and long-term (L) acclimation experiments sampled across temperature treatments (27°C control, 29.5°C, 32°C, and 34.5°C heat stress). In this dataset, T1 corresponds to the acute heat stress phase in the experiment.

Paired-end raw reads were trimmed and filtered for low-quality bases using Trimmomatic v0.39^25^. Quality assessment was performed with FastQC ^26^ and MultiQC^27^. Filtered reads were aligned to the concatenated *S. pistillata*–*Symbiodinium microadriaticum* reference genome using HISAT2 2.2.1^17^. Gene-level read counts were generated using featureCounts 2.1.1^28^ based on the extended gene annotation. Low-quality libraries were removed based on number of reads (less than 1 million reads) remaining after quality filtration steps as well as their mapping rates (less than 9%). The retained library sizes ranged from 1.68 million to 114.17 million reads, with a median at 23.03 million. Read alignment to the concatenated reference varied across samples from 9.85% to 91.44% (median = 75.87%; Supplementary Table S1). Gene-assignment rates from featureCounts ranged from 3.5% to 60.9% (median = 40%; Supplementary Table S1). Genes with row sums < 10 across all samples were removed to reduce noise and improve statistical power. Principal component analysis was conducted on normalized counts, and PCA plots were generated using ggplot2 3.5.2^29^ to compare global expression patterns with those reported in the original publications.

Differential expression analysis was performed using DESeq2 1.46.0^30^. Pairwise comparisons between experimental conditions were conducted within each dataset. Genes were considered differentially expressed if they met the criteria |log2FC| > 1 and adjusted p-value < 0.05. Differentially expressed genes were identified for all relevant condition contrasts within each dataset. For downstream analyses, two DEG sets were selected: ICN 36 vs 30, representing the transcriptional response of the most heat-sensitive population (ICN) under extreme heat stress (36°C vs 30°C) from Voolstra 2021, and short-term 34.5 vs 27 at T1, representing the short-term acute heat stress response at timepoint T1 (34.5°C vs 27°C), which exhibited the strongest transcriptional response from Savary 2021. For visualization only, TMM normalization was calculated using edgeR 4.4.0^31^ to account for differences in library composition across samples, and TMM-normalized expression values were used in heatmaps.

### 6. Intersection Analysis Between bRNAseq DEGs and scRNA Neuronal Gene Sets

To identify neuron-enriched genes responsive to heat stress, neuron-specific gene lists derived from single-cell analysis were intersected with heat stress DEG lists obtained from bulk RNA-seq analyses. Four pairwise intersections were performed, combining two neuron-enriched gene sets (*neuron_highexpr* and *neuron_predominant*) with two bulk heat stress DEG sets (ICN 36 vs 30 and short term 34.5 vs 27 at T1). Gene lists were matched by gene ID, and only genes present in both corresponding input lists were retained in each intersection. The resulting intersected gene sets represent candidate neuron-enriched genes associated with heat stress in bulk tissue experiments.

### 7. Cell-Type Deconvolution of Bulk RNAseq Data

To estimate cell-type-specific (CTS) gene expression profiles from bulk RNA-seq data and provide a complementary estimate of neuron-associated expression changes, computational deconvolution was performed using InstaPrism 0.1.6^32^. Deconvolution leverages single-cell reference expression profiles to infer the proportional contribution of each cell type within bulk samples and to estimate CTS gene expression levels from mixed tissue data. The annotated Seurat object from scRNA-seq analysis and expression matrices from bulk RNA-seq were used as inputs to InstaPrism. Deconvolution was performed using the InstaPrism function with default reference-preparation and outlier-filtering parameters. InstaPrism generated (i) a theta matrix representing estimated cell type fractions for each bulk sample, and (ii) a three-dimensional gene expression array (Z array; samples × genes × cell types) providing inferred CTS expression estimates for each gene in each sample.

For neuronal expression analysis, the inferred neuronal expression matrix was extracted from the Z array by selecting the neuron cluster across all samples and genes. To reduce noise from minimally expressed genes in the deconvolved data, genes with row sums ≤ 1 across all samples were removed. Differential expression analysis was performed using Mann–Whitney U tests comparing control and heat-stressed samples within each population or experimental condition. P values were adjusted across genes within each comparison using the Benjamini–Hochberg false-discovery-rate. Genes were classified as significantly differentially expressed using the criteria |log2 FC| > 1 and adjusted p-value < 0.05. DEG analyses were conducted separately for each population in the Voolstra et al (2021) dataset (ICN, AFR, ETR, PTR) and for each experimental condition in the Savary et al. (2021) dataset (both short term and long term at T1), using extreme temperature contrasts (36°C vs 30°C in Voolstra; 34.5°C vs 27°C in Savary). For downstream analyses, only differentially expressed genes from ICN 36 vs 30 and from the short-term T1 comparison 34.5 vs 27 were retained, as these comparisons exhibited the strongest transcriptional responses to heat stress.

### 8. Consensus Candidate Gene Identification and Visualization

To identify consensus heat-associated neuronal candidate genes, results from two complementary analytical routes were compared: (i) intersection of bulk DEGs with single-cell neuronal gene sets and (ii) deconvolution-based neuronal DEG analysis. Comparisons were performed across two independent datasets (Voolstra et al 2021^23^, ICN 36 vs 30; Savary et al 2021^24^, short-term 34 vs 27 at T1) and two neuron gene set definitions (*neuron_highexpr* and *neuron_predominant*). For each dataset and gene set combination, genes were matched by gene ID between the two analytical methods. Only genes showing consistent regulatory direction (consistently upregulated or downregulated under heat stress) were retained. High-confidence candidates were required to be detected by both analytical routes, responsive in both independent studies, supported by both neuronal gene set definitions, and consistently regulated in the same direction across all comparisons.

Expression patterns of the final candidate genes were visualized. In single-cell data, candidate gene expression across cells and clusters was visualized using FeaturePlot and DotPlot in Seurat 5.3.0^20^, and in R 4.4.2^19^ with ggplot2 3.5.2^29^. In bulk RNA-seq data, gene expression was visualized using heatmaps generated in R 4.4.2^19^ with ggplot2 3.5.2^29^, grouped by population or condition and temperature. Inferred neuronal expression estimates derived from InstaPrism 0.1.6^32^ deconvolution were visualized using the same grouping framework.

### 9. Targeted sequence-based characterization of consensus candidate proteins

Predicted protein sequences for the nine consensus candidates were individually evaluated using complementary sequence-based approaches, depending on the predicted protein features and initial annotation. Sequence similarity was assessed using BLASTp against the NCBI non-redundant (nr) protein database^33^, and conserved domains and protein families were evaluated using NCBI CD-Search^34^ (CDD; E-value cutoff 0.01) and InterPro^35^. SignalP 6.0^36^ (Eukarya, slow mode) was used to evaluate potential N-terminal signal peptides in g15744 and g29914, and DeepTMHMM 1.0^37^ was used to evaluate transmembrane topology in g1393, g5684, g15744, g25745, and g29914. Potential phosphorylation sites in g21193 were predicted using NetPhos 3.1 b^38^ (scores >0.5). Additional analyses were performed on selected candidates: For phylogenetic analysis of the TRPA-family candidate g25745, TRPA protein sequences from model species, including *Homo sapiens*, *Mus musculus*, and *Drosophila melanogaster*, were retrieved from NCBI. We included all proteins annotated as TRPA in the *Stylophora* genome, together with hits identified by searching its predicted proteome with g25745 using BLASTp (E-value cutoff: 1 × 10□³□). Protein sequences were aligned with MAFFT^39^ and trimmed with trimAl^40^. A maximum-likelihood tree was reconstructed in IQ-TREE 2^41^ using a substitution model selected by ModelFinder and 1,000 ultrafast bootstrap replicates, then visualized in R with phytools. In addition, the Class A/rhodopsin-like GPCR g5684 was compared with experimentally deorphanized *Nematostella vectensis* GPCRs^42^ using local pairwise protein-sequence alignments implemented with the Bio.Align.PairwiseAligner module in Biopython^43^, with the BLOSUM62 substitution matrix and gap-opening and gap-extension penalties of -11 and -1, respectively.

### 10. Generative AI Use Statement

Initial protein-sequence interpretation and integration of results from complementary annotation resources were assisted by ChatGPT (GPT-5.6 Sol; OpenAI; accessed September 2026). All candidate annotations proposed during this process were subsequently evaluated by the authors through manual inspection of the protein sequences and human curation of the underlying sequence-similarity, conserved-domain, topology, and phylogenetic evidence. AI-generated interpretations were not accepted as annotation evidence independently of these analyses, and final annotations were assigned conservatively at the most specific level supported by concordant evidence. Codex (OpenAI) assisted with writing and revising analysis code. The authors specified the key analytical parameters and reviewed AI-assisted code before use. ChatGPT was also used to improve human-generated text for readability and style. All scientific interpretations and final manuscript content were reviewed and approved by the authors.

## Results

### 1. scRNA-seq Mapping and Clustering

Extension of annotated gene intervals increased gene model length. A total of 18,387 of 30,756 genes (59.8%) were extended at the 3’ end. The average gene length increased by 523 bp, from 5,915 bp in the original annotation to 6,438 bp after extension. The extended gene model was used for subsequent scRNAseq and bulk RNAseq analyses. scRNA mapping recovered 18,459 cells. The median number of detected genes per cell was 744. The overall alignment rate to the concatenated host-symbiont genome was 66.8%. Of all high-quality reads, 53.3% mapped to the genome and 43.2% mapped to annotated *S. pistillata* genes. A total of 25,048 genes were detected across all cells. Valid barcodes comprised 96.1% of reads, and 100% of UMI sequences were valid. Sequencing saturation reached 87.8%, and Q30 scores exceeded 93% for RNA reads. Graph-based clustering of filtered cells using the first 20 principal components identified 16 distinct clusters (Supplementary Figure S1). The top 20 marker genes were identified for each cluster, yielding 320 marker genes in total, of which 95 (29.69%) had functional annotations with assigned gene names (Supplementary Table S2).

### 2. Annotation of scRNA Cell Clusters

To annotate cell clusters, the top 25 marker genes from 12 previously defined cell types (N=300) in Levy et al. (2021) were retrieved and mapped to the proteome inferred from the chromosome-level genome assembly using OrthoFinder (Supplementary Table S3). Among the 300 reference marker genes, 169 (56.3%) had identifiable orthologs in the chromosome-level genome. Of these, 156 genes exhibited a one-to-one orthologous relationship, indicating direct correspondence between gene models across genome versions (Supplementary Table S3).

Expression of mapped orthologs was visualized across cell clusters using dot plots and UMAP feature plots (Supplementary Files in Figshare Repo). Neuronal markers are shown as an example. Multiple neuronal orthologs, including g20052 (PKD2), g22183 (CACNA1G), g9687 (CACNA1E), and g17683 (UTRN), showed enriched expression in discrete clusters. Clusters 6, 11, 13, and 15 displayed the strongest neuronal gene signatures and were designated as neuronal cell populations (Supplementary Figure S1). In parallel, a keyword-based search strategy was applied using curated functional terms associated with 12 general cell types from Levy et al. (2021) (Supplementary Table S4, S5). The number of predominant gene hits ranged from 8 (germline-associated keywords) to 472 (gland-associated keywords). Dot plots revealed that many keyword-derived genes showed weak expression patterns. However, some genes exhibited strong, cluster-restricted expression. Genes associated with symbiont metabolism and host-symbiont interaction, including g15457 (UGP2), g14213 (CHAC1), and g9952 (GALT), showed strong expression in cluster 4 (Supplementary Files in Figshare Repo).

### 3. Identification of Neuronal Genes

Differential expression analysis comparing neuronal cells to all other cells identified 384 genes that were highly expressed in neurons (log2fc ≥ 1.0 and Padj < 0.05; *neuron_highexpr;* Supplementary Table S7). Of these, 203 genes (52.86%) lacked functional annotation from emapper. A complementary proportional expression criterion identified 283 genes for which at least 80% of counts were assigned to neuronal cell clusters (*neuron_predominant;* Supplementary Table S8). Among these, 217 genes (76.68%) were unannotated. Intersection of *neuron_highexpr* and *neuron_predominant* yielded 83 overlapping genes (Fig. 2, Supplementary Table S9). Of these, 62 genes (74.7%) lacked functional annotation.

**Figure 2.**
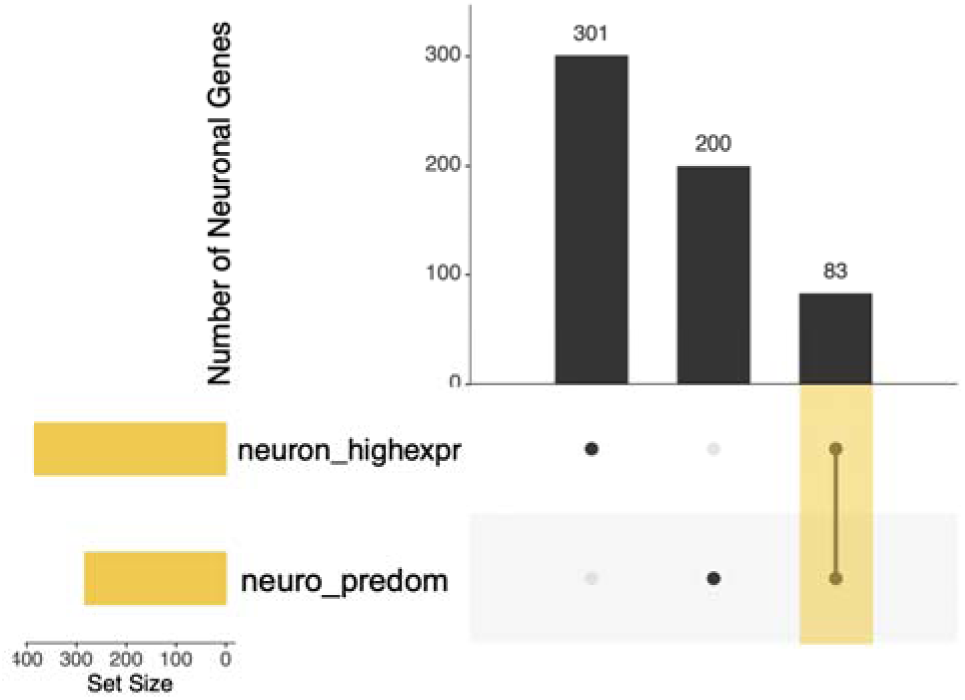
Identification of neuronal genes. Neuronal genes were defined using two complementary criteria: The *neuron_highexpr* set includes genes with significantly higher expression in neuronal cells compared to all other cell types. The *neuron_predominant* set includes genes with expression predominantly restricted to neuronal cells. Intersection of these two sets identifies 83 genes that are both strongly enriched and specifically expressed in neurons

Among the annotated genes, several showed sequence similarities to genes with established neuronal functions in other animals, including NPTX1-like neuronal pentraxin, OPN4-like opsin, CACNA1C-, CACNA1G- and KCNA2-like voltage-gated ion channels, and a TRPA1-like ion channel. Importantly, these annotations are based on sequence homology and do not necessarily establish equivalent functions in corals. The recovery of genes associated with neuronal signaling and sensory functions was consistent with the neuronal identity of these gene sets.

### 4. Differential Gene Expression Analyses of Bulk RNA-seq

The largest transcriptional response to heat stress was observed in response to the most extreme temperature exposures, particularly in the ICN population that originated from the colder northern Red Sea in Gulf of Aqaba. The ICN 36 vs 30 comparison identified 8,818 DEGs, consistent with the strongest response previously reported by Voolstra et al. (2021) (Supplementary Table S10). This number was approximately ten-fold greater than that observed in other populations (Supplementary Figure S2). Within the ICN population, the 36°C vs 33°C comparison identified 6,867 DEGs. The majority of these genes were downregulated (Supplementary Figure S2). Across populations, larger temperature contrasts consistently produced more extensive transcriptional responses (Supplementary Figure S2). For example, 36°C versus 30°C contrasts yielded substantially more DEGs than 33°C versus 30°C comparisons (Supplementary Table S10).

In the Savary et al. (2021) dataset, short-term acute heat stress led to stronger transcriptional changes than long-term thermal acclimation. The short term 34.5 vs 27 comparison at T1 identified 7,761 DEGs, whereas the long term 34.5 vs 27 comparison at T1 identified 2,718 DEGs (Supplementary Table S10). Like the results from Voolstra 2021 dataset, 34.5 vs 27 contrasts produced notably more DEGs than 32 vs 27 contrasts (Supplementary Figure S3). Overlap between long-term and short-term responses was limited, with fewer than 10% of DEGs shared between the two comparisons (Supplementary Figure S3).

Cross-study comparison of the extreme heat contrasts revealed substantial overlap between the ICN 36 vs 30 and short-term 34.5 vs 27 at T1. UpSet analysis identified 2,472 shared downregulated genes and 817 shared upregulated genes between these contrasts (Supplementary Figure S4). This cross-study concordance provided the basis for focusing subsequent analyses on genes supported by both independent datasets.

### 5. Intersection of Bulk Heat Stress DEGs with Neuronal Candidate Genes

Among the 384 *neuron_highexpr* genes, 138 (35.9%) overlapped with DEGs from the ICN 36 vs 30 comparison and 153 (39.8%) overlapped with DEGs from the short-term 34.5 vs 27 at T1 comparison. Of these, 75 genes were shared between the two bulk contrasts (Supplementary Figure S5). Among the 283 *neuron_predominant* genes, 53 (18.7%) overlapped with ICN 36 vs 30 DEGs and 43 (15.2%) overlapped with short-term 34.5 vs 27 at T1 DEGs, with 21 genes shared between the two bulk contrasts (Supplementary Figure S5).

In total, 216 of 384 *neuron_highexpr* genes (56.2%) and 75 of 283 *neuron_predominant* genes (26.5%) responded to heat stress in at least one bRNAseq dataset (Supplementary Figure S5). Conversely, neuronal genes represented a relatively small fraction of all heat-responsive genes. Of the 8,818 DEGs identified in the ICN 36 vs 30 comparison, 138 (1.6%) overlapped with *neuron_highexpr* and 53 (0.6%) with *neuron_predominant*. Of the 7,761 DEGs identified in the short-term 34.5 vs 27 at T1 comparison, 153 (2.0%) overlapped with *neuron_highexpr* and 43 (0.6%) with *neuron_predominant*.

Fourteen genes were shared across all four gene sets (*neuron_highexpr*, *neuron_prodominant*, ICN 36 vs 30, and short-term 34.5 vs 27 at T1; Figure 2; Supplementary Table S11). Of these, four lacked functional annotation. Homology-based annotations for the remaining genes included CALM1-like, SPRYD3-like, PKNL-like, ADRB2-like, RET-like, TRPA1-like, and SKP2-like genes (Supplementary Table S11). Thus, intersection of the single-cell neuronal gene sets with the two independent bRNAseq datasets reduced the broader transcriptional response to a small set of neuron-enriched genes consistently associated with heat stress.

### 6. Deconvolution-based Neuronal Differential Expression under Heat Stress

As a complementary approach to the gene-set intersection analysis, cell-type deconvolution was used to estimate cell-type fractions and infer cell-type-specific (CTS) expression profiles from the bRNAseq data. Whereas the intersection analysis evaluated heat responsiveness among genes predefined as neuron-enriched in the single-cell atlas, deconvolution provided neuronal expression estimates for the broader set of genes represented in the reference.

Estimated neuron cell fractions did not differ significantly between control and heat-stress conditions, consistent with the relative stability of neuronal populations under heat stress in a single-cell atlas in corals^44^. Estimated fractions of several other cell types differed between conditions. In the ICN population, estimated fractions of calicoblasts, epidermal cells, alga-hosting cells, and gland cells decreased, whereas the gastrodermal fraction increased at 36°C (adjusted p < 0.05; Supplementary table S12, S13).

In the Voolstra (2021) dataset, deconvolution of the ICN 36 vs 30 comparison identified 6,441 significant neuronal DEGs, of which 99.7% were downregulated. In contrast, no significant neuronal DEGs were detected in the other source reef populations under the same thermal contrast (AFR, ETR, or PTR; 36°C vs 30°C). In the Savary (2021) dataset, short-term heat exposure between 34.5°C and 27°C at T1 identified 3,079 significant neuronal DEGs, of which 88.9% were downregulated. No significant neuronal DEGs were detected under long-term exposure (long-term 34.5 vs 27 at T1).

These results indicate population- and exposure-dependent neuronal transcriptional responses to heat stress, with extensive downregulation of neuron-associated gene expression in the ICN population and during short-term acute heat exposure from the two datasets respectively.

### 7. Cross-Study Consensus Prioritizes Neuronal Heat-stress Genes

We then applied a stringent consensus criterion requiring genes to (1) satisfy both single-cell based neuronal gene definitions *(neuron_highexpr and neuron_predominant)*, (2) be differentially expressed in both independent bRNAseq studies, (3) be identified as neuronal DEGs by deconvolution in both studies, and (4) show the same direction of differential expression in the two extreme contrasts. This analysis identified nine consensus candidate genes, all of which were downregulated in both studies (Fig. 3, Table 1): g1393, g5684, g7849, g15744, g21193, g23815, g24582, g25745, and g29914.

**Figure 3.**
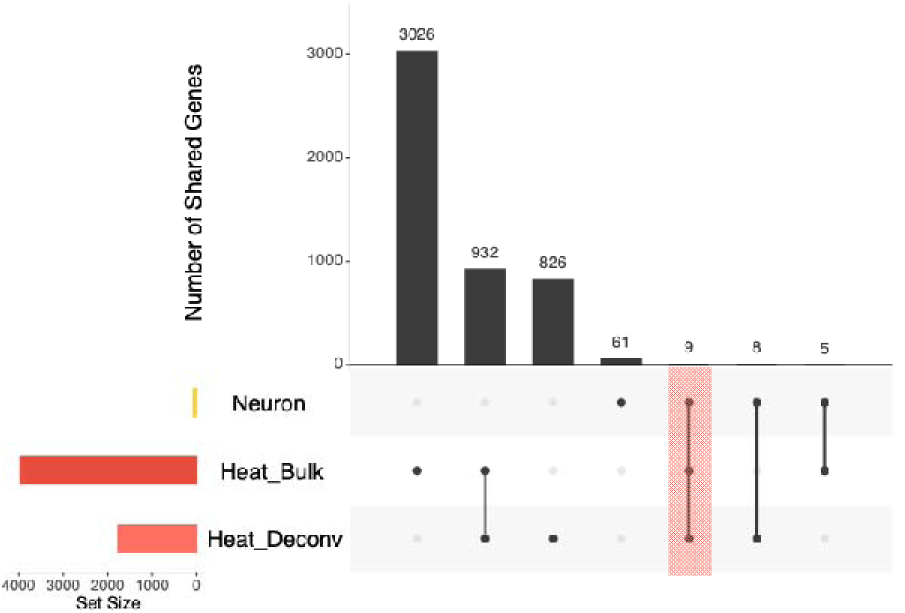
Overlap of neuronal and heat responsive gene sets. Intersection of neuronal genes, bulk heat responsive genes, and deconvolution-derived heat responsive genes in neurons identifies 9 high-confidence candidate genes responsive to heat stress in neurons.

**Table 1.** Initial annotations and revised sequence-based characterization of the nine consensus neuronal heat-stress candidate genes. GPCR, G protein-coupled receptor; aa, amino acids. Initial annotations represent annotations available before targeted sequence-based characterization. Revised characterizations were assigned conservatively based on sequence similarity, predicted protein architecture, conserved domains, topology, and phylogenetic evidence, as applicable.

| Gene ID | Initial annotation | Revised characterization | Key evidence / interpretation |
| --- | --- | --- | --- |
| g1393 | SPRYD3-like; TRPC4-like | Multipass membrane protein; specific family unresolved | Predicted protein architecture does not convincingly support a canonical TRPC4 channel and raises the possibility of a chimeric or misassembled gene model |
| g5684 | ADRB2-like | Partial/truncated Class A/rhodopsin-like GPCR; receptor identity unresolved | GPCR architecture supported, but ADRB2 assignment is not. Sequence similarity to experimentally deorphanized *Nematostella* amidated-neuropeptide-responsive GPCRs suggests possible peptidergic function but does not establish ligand specificity |
| g7849 | Unannotated | Uncharacterized cysteine-rich protein | Cysteine-rich architecture; protein family and molecular function unresolved |
| g15744 | Unannotated | Predicted secreted protein belonging to a conserved coral protein family of unknown function | DeepTMHMM predicts an N-terminal signal peptide (aa 1–17) and no transmembrane regions; BLASTp identifies full-length homologs across multiple coral genera |
| g21193 | Unannotated | Highly basic intracellular protein; function unresolved | Multiple predicted phosphorylation sites, including numerous putative PKC phosphorylation sites |
| g23815 | Unannotated | Calmodulin/centrin-family EF-hand Ca<sup>2+</sup>-binding protein; specific paralog unresolved | Multiple EF-hand Ca<sup>2+</sup>-binding domains support family-level assignment |
| g24582 | CALM1-like | Calmodulin/centrin-family EF-hand Ca<sup>2+</sup>-binding protein; specific paralog unresolved | Multiple EF-hand Ca<sup>2+</sup>-binding domains support family-level assignment; specific CALM1 orthology cannot be resolved |
| g25745 | TRPA1-like | TRPA-family ion channel | Ankyrin-repeat-rich N-terminal region and C-terminal transmembrane channel domain; phylogenetic placement within a diversified group of coral TRPA-family channels does not support simple one-to-one orthology with bilaterian TRPA1 |
| g29914 | Unannotated | Predicted secreted protein belonging to a conserved coral protein family of unknown function | DeepTMHMM predicts an N-terminal signal peptide (aa 1–35) and no transmembrane regions; BLASTp identifies full-length homologs across multiple coral genera |

Initial automated annotations assigned putative identities to four candidates: g1393 (SPRYD3-like), g5684 (ADRB2-like), g24582 (CALM1-like), and g25745 (TRPA1-like). The remaining five lacked informative functional annotations. Because homology-based annotations can be particularly uncertain in evolutionarily divergent non-model organisms, we subsequently performed targeted sequence-based characterization of all nine candidate proteins (Section 8). Visualization of the scRNA-seq data showed neuronal enrichment of all nine candidates (Fig. 4). Their expression patterns in the bRNAseq datasets showed consistent downregulation in the two extreme contrasts used for consensus prioritization, although expression patterns varied across other populations, temperatures, and exposure durations (Fig. 5). Eight of the nine consensus candidates showed their highest mean expression in neuronal cluster 6 (Supplementary Figure S6). The remaining candidate, g7849, was expressed in cluster 6 but showed its highest mean expression in neuronal cluster 11 (Supplementary Figure S6). Together, these analyses prioritized nine neuronal genes supported by two independent analytical approaches and two independent heat-stress datasets.

**Figure 4.**
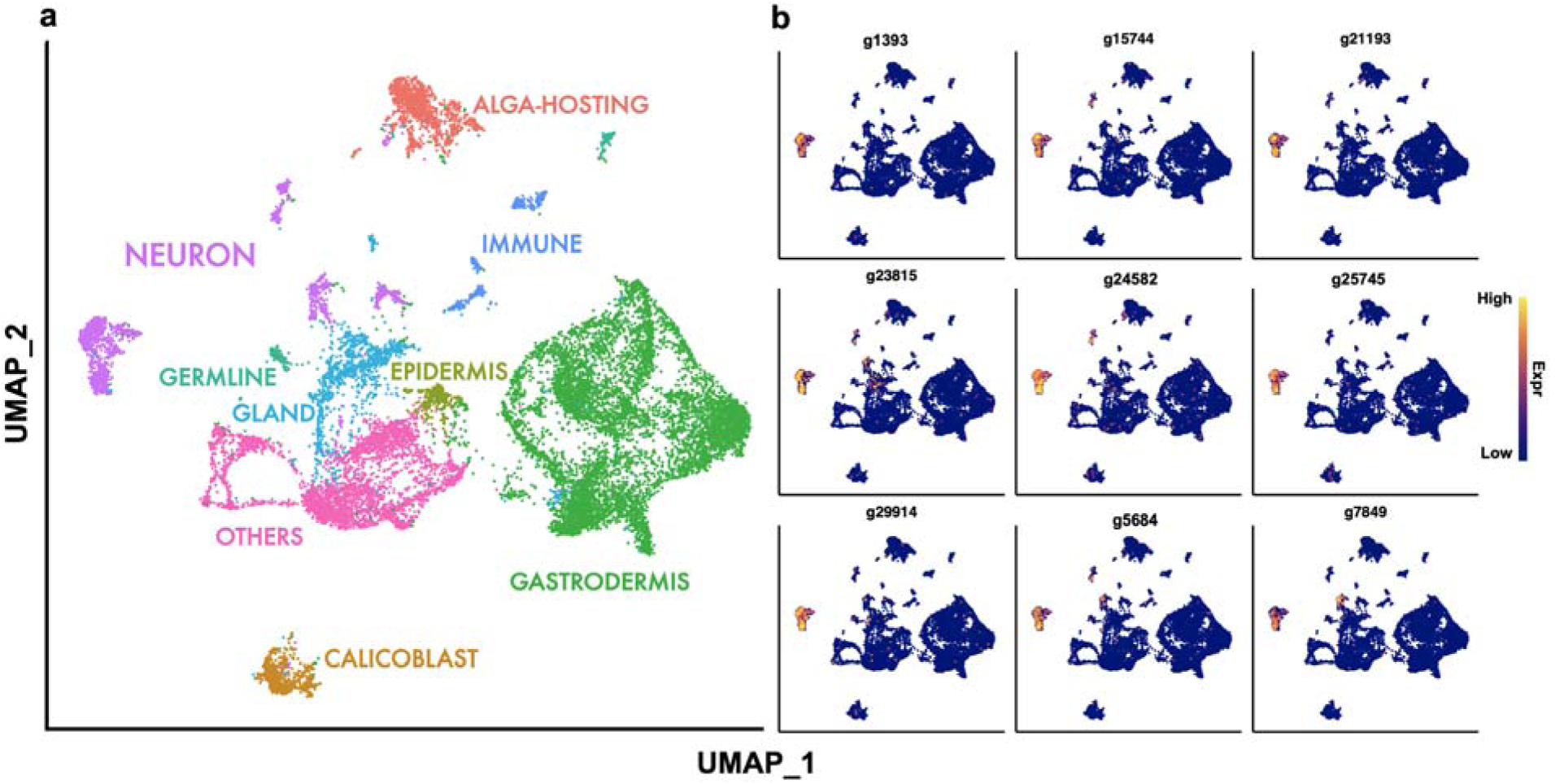
Cell and expression patterns of candidate neuronal heat responsive genes. a) UMAP visualization of scRNA sequencing data showing clustering and annotation of major cell types; b) Feature plots showing expression of nine candidate genes across all cells. Color scale indicates expression level, with dark blue representing no expression and yellow representing high expression.

**Figure 5.**
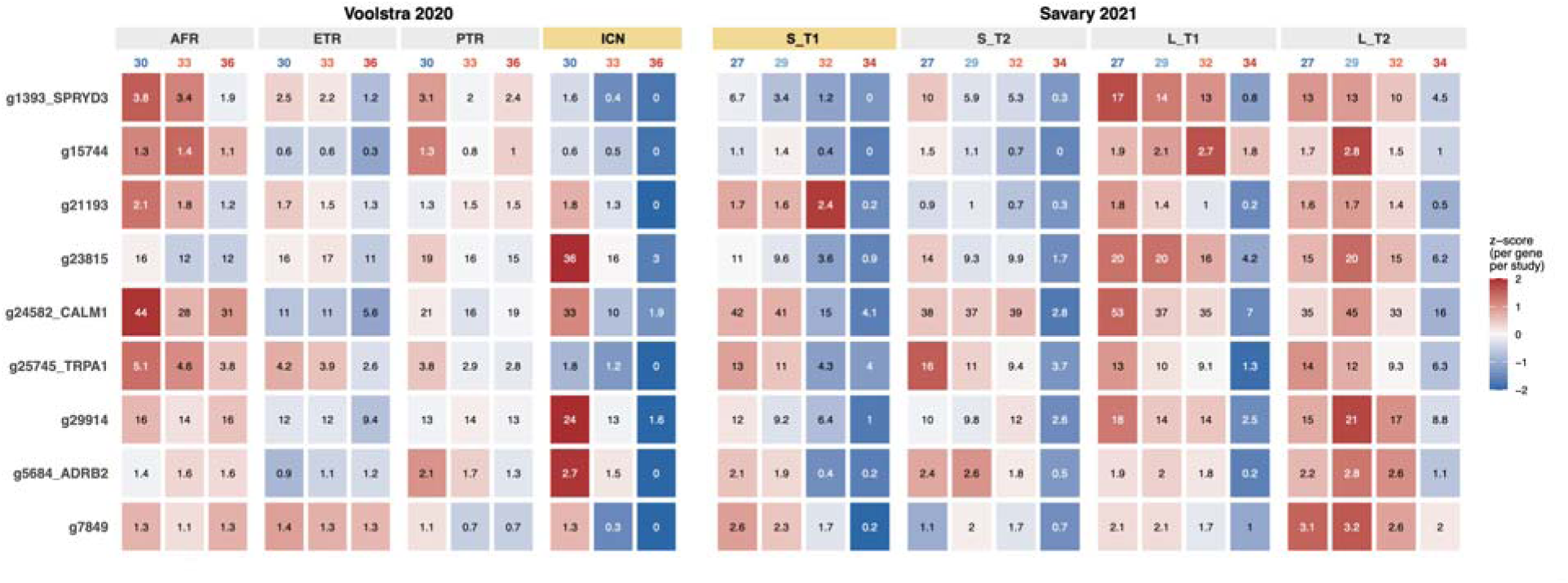
Expression patterns of candidate neuronal genes across heat stress conditions. Heatmap showing expression of nine candidate genes across temperature conditions and treatment groups in Voolstra 2021 and Savary 2021 datasets. Values are shown as average expression in TMM within each group.

### 8. Targeted sequence-based characterization of consensus candidate proteins

Because several consensus candidates were either unannotated or had automated annotations based primarily on homology to distantly related organisms, we performed targeted sequence-based characterization of all nine predicted proteins (Table 1). This analysis provided varying levels of support for their initial annotations and, in some cases, supported revised or additional characterization. The predicted g25745 protein displayed the characteristic architecture of a TRPA-family ion channel, including an extended N-terminal ankyrin-repeat-rich region and a C-terminal transmembrane channel domain. Phylogenetic analysis placed g25745 within a diversified group of coral TRPA-family channels, supporting its identification as a TRPA-like channel but not a simple one-to-one orthology with bilaterian TRPA1 (Supplementary Figure S7). The previously unannotated g23815 and CALM1-like g24582 contained multiple EF-hand Ca²□-binding domains consistent with calmodulin/centrin-family proteins, although their specific paralog identities could not be resolved.

Other automated annotations were less well supported. g1393 was initially annotated as SPRYD3-like, while additional sequence-similarity searches suggested similarity to TRPC4. However, its predicted protein architecture did not convincingly support a canonical TRPC4 channel and raised the possibility of a chimeric or misassembled gene model. g5684 contained sequence features consistent with a Class A/rhodopsin-like GPCR, but sequence similarity was insufficient to support its automated ADRB2 annotation or a specific receptor identity. Comparisons with experimentally characterized cnidarian GPCRs revealed greatest similarity to amidated-neuropeptide-responsive GPCRs from *Nematostella vectensis*, although this was insufficient to assign a specific ligand or receptor subtype.

The remaining candidates lacked sufficiently strong homology for family-level functional assignments. Sequence analysis nevertheless identified two predicted secreted proteins (g15744 and g29914) with strong sequence conservation among uncharacterized coral proteins, indicating that each belongs to a conserved coral protein family of unknown function. Additional features included a cysteine-rich protein (g7849), and a highly basic intracellular protein with multiple predicted phosphorylation sites, including numerous putative PKC phosphorylation sites (g21193). These candidates were therefore retained with conservative descriptions based on the available evidence (Table 1).

## Discussion

Cell-type resolution remains a major limitation in environmental transcriptomics, particularly for non-model organisms in which experimental single-cell datasets across different treatments are still sparse. Here, we addressed this challenge by using an existing *Stylophora pistillata* single-cell atlas to add cellular context to two independently generated bulk heat-stress datasets. By integrating neuronal gene enrichment, cross-study differential-expression overlap, and deconvolution, we narrowed thousands of heat-responsive genes to nine neuron-enriched candidates consistently supported across both studies and analytical approaches. Because cell-type association provides information independent of functional annotation, this framework can prioritize poorly characterized genes that pathway-based analyses would overlook. All nine were downregulated under the heat-stress conditions used for consensus prioritization. These included a TRPA-family ion channel, two calmodulin/centrin-family Ca^2+^-binding proteins, a Class A/rhodopsin-like GPCR, and several proteins whose functions remain unresolved. Their coordinated downregulation identifies a reproducible component of the neuronal response to heat stress, and these candidates provide a tractable starting point for spatial localization (e.g., HCR-FISH, immunohistochemistry) and targeted functional perturbation (e.g., CRISPR). At the same time, the uncertain functional annotation of several candidates highlights the challenges of inferring gene function from homology in non-model organisms.

### A conservative framework for prioritizing cell-type-associated environmental responses

This study presents a reproducible workflow for prioritizing cell-type-associated environmental-response genes by integrating complementary single-cell and bulk transcriptomic datasets. The single-cell reference was critical for providing cell-type context, but represented only a single environmental condition. In contrast, the bRNA-seq studies examined expression across multiple thermal contrasts, but samples comprised heterogeneous coral tissues. Integrating these datasets allowed us to associate heat-responsive genes with neuronal expression without requiring matched single-cell sequencing for each experimental condition. Our approach therefore prioritizes robustness over completeness, with the goal of identifying a tractable set of candidates for further investigation rather than reconstructing the complete neuronal heat-stress transcriptome.

To reduce uncertainty, we adopted a conservative strategy for defining neuronal genes, focusing on those strongly enriched or predominantly expressed in neuronal clusters. Genes shared between neurons and other cell types were therefore not systematically captured, even though changes in their expression could contribute meaningfully to the overall neuronal response^45^. This restriction was intended to minimize false-positive assignments at the cost of excluding potentially relevant genes. The thresholds used here are not intrinsic to the framework and could be adjusted depending on whether a particular application prioritizes sensitivity or specificity.

Deconvolution uses a single-cell reference to infer cell-type-associated signals from mixed bulk samples. A critical assumption of this approach is that cell-type-specific expression remains stable across conditions^46^. Consequently, the approach is better suited to identifying heat-responsive genes that are already expressed in neurons under control conditions than genes that are induced specifically by heat. This asymmetry is particularly relevant because all nine consensus candidates identified here were downregulated under the two heat-stress conditions used for prioritization and therefore should not be interpreted as evidence that the neuronal response to heat is predominantly downregulatory. Although estimated^44^, changes in expression of neuronal markers can complicate inference of cell abundace from bulk transcriptomes without paired single cell libraries. The deconvolution analysis was therefore used as a complementary approach for candidate prioritization rather than as a direct measurement of neuronal abundance or a comprehensive reconstruction of neuronal transcriptional dynamics.

A major advantage of this approach is its ability to reduce the complexity inherent in environmental transcriptomic datasets. Bulk RNAseq experiments commonly identify hundreds or thousands of differentially expressed genes, while functional validation remains a major bottleneck in linking genotype to phenotype. This problem is amplified in non-model organisms, where functional annotation is often incomplete and pathway-based analyses can overlook genes without established homologs or functions. By incorporating cell-type association as an additional source of biological information alongside functional annotation, our framework can prioritize both well-annotated and poorly characterized genes for further investigation. The resulting nine neuronal candidates provide a tractable starting point for downstream approaches such as spatial localization (e.g., immunohistochemistry, HCR-FISH) and targeted functional perturbation (e.g., CRISPR).

Although demonstrated here using neuronal responses to heat stress, the framework is not restricted to either neurons or temperature. The same strategy could be applied to other cell populations and environmental responses, such as calcifying cells in studies of biomineralization or gastrodermal cells in studies of symbiosis. More broadly, it provides a way to leverage existing single-cell atlases to extract cell-type-associated information from independently generated environmental transcriptomic datasets.

### Candidate genes provide insights into coral neuronal signaling and heat response

Like all animals, corals must continuously sense and integrate information from the environment and internal physiological state. Environmental cues include light, temperature, oxygen, pH and other chemical signals, as well as mechanical stimuli. These signals influence fundamental aspects of coral biology, including feeding, defensive responses, tissue contraction, reproduction, tissue regeneration, and growth^50^. Temperature presents a particularly important challenge because it strongly influences metabolic rate, membrane properties, and cellular homeostasis, all of which are critical for neuronal function. Maintaining function across changing thermal conditions therefore likely requires both sensing temperature and adjusting neuronal and physiological processes accordingly^51^. The nervous system may therefore help integrate environmental information and coordinate organismal responses to changing conditions^52^. Under more extreme or prolonged thermal stress, disruption of neuronal function could compromise this coordination, potentially amplifying the effects of stress across multiple physiological processes.

Unlike the nervous system of bilaterian animals, which is typically organized around centralized structures such as brains and ganglia, cnidarian nervous systems consist of neurons distributed throughout the animal without centralized nervous organs^53^. This organization makes understanding nervous system function at the cellular level particularly important. Identifying different neuronal cell types, the genes they express, and how their molecular profiles change with environmental conditions provides a pathway to understanding how the coral nervous system senses information and coordinates physiological responses. Single-cell transcriptomics is beginning to provide this resolution, revealing molecularly distinct neuronal populations in corals and other cnidarians^9,54–57^.

However, the physiological functions of these neuronal populations and much of their underlying molecular machinery remain poorly characterized. Some components likely reflect deeply conserved features of animal neurobiology (e.g. ion channels, receptors, intracellular signaling proteins), whereas others may be highly divergent or specific to cnidarians or reef-building corals, making their functions difficult to infer from conventional model organisms. This incomplete knowledge complicates the interpretation of neuronal responses to environmental stress but also makes cell-type-resolved environmental transcriptomics particularly informative. By identifying genes that are enriched in neuronal cells and consistently respond to thermal stress, our approach provides experimentally testable candidates for both conserved and previously uncharacterized components of coral neuronal function.

Among the better characterized candidates, g25745 was identified as a TRPA1-like channel. TRPA channels are polymodal sensory channels that can respond to chemical, mechanical, and thermal stimuli, although their sensory properties vary substantially among lineages^58,59^. The predicted g25745 protein contains an extended ankyrin-repeat-rich N-terminal region and a C-terminal transmembrane channel domain characteristic of TRPA-family channels. Phylogenetic analysis further places g25745 within a large expansion of coral TRPA-family channels and does not support simple one-to-one orthology with bilaterian TRPA1. Without one-to-one orthology, the sensory function of g25745 cannot be inferred directly from bilaterian TRPA1. ^60^ The TRPA-like identity of g25745, together with its neuronal enrichment and consistent downregulation during heat stress, makes it a particularly interesting candidate for environmental sensing. However, whether it directly senses temperature, or even whether temperature is its primary stimulus requires experimental validation.

Evidence from other anthozoans also links TRPA-family channels with responses to thermally variable environments. In *Acropora tenuis* from extreme environments in Bouraké lagoon, a TRPA1-family gene showed a strong signature of environmental selection, including multiple environmentally associated variants and a nonsynonymous variant predicted to affect protein structure^61^. However, because temperature, pH, oxygen, and other environmental variables covary in this habitat, this signal cannot be attributed exclusively to temperature. In the anemone *Exaiptasia diaphana*, TRPA1-immunoreactive protein abundance increased under severe thermal stress^62^, in contrast to the downregulation of g25745 identified in our heat-stress datasets. Moreover, TRPA1 immunoreactivity in *Exaiptasia* was reported primarily in epidermal and gastrodermal cells that did not appear to be neuronal, suggesting that TRPA-family proteins may participate in diverse heat-associated responses in multiple anthozoan cell types. Differences in species, cell type, experimental conditions, and protein versus transcript measurements preclude direct comparison of the direction of these responses. Together, these observations support further investigation of anthozoan TRPA-family channels in environmental responses, while providing no basis for assuming a conserved cellular role or sensory function.

Several of the other candidates illustrate how functional interpretation in corals can be misled by automated annotations that imply more specificity than the sequence supports. For example, g1393 was initially annotated as SPRYD3-like and then proposed to bear similarity with TRPC4. This was an intriguing possibility because TRPC channels have been implicated in temperature-sensitive signaling^63^. However, closer examination of g1393 did not convincingly support a canonical TRPC4 channel and instead raised the possibility of a chimeric or misassembled gene model. Similarly, g5684 was initially annotated as ADRB2-like, despite phylogenetic evidence indicating that canonical adrenergic receptor families diversified within Bilateralia^64^. Sequence analysis instead supported a Class A/rhodopsin-like GPCR, with greatest similarity to experimentally deorphanized *Nematostella* amidated-neuropeptide-responsive GPCRs. Finally, g23815 and g24582 were identified as calmodulin/centrin family containing multiple EF-hand Ca^2+^ binding domains, although their specific paralog identities could not be resolved. Together, these examples illustrate both the difficulty of assigning specific molecular functions from homology across deep evolutionary distances and the value of sequence-based characterization for refining automated annotations.

Eight of the nine candidates showed their highest mean expression in neuronal cluster 6. Importantly, cluster identity was not used to select the consensus candidates, so their convergence on cluster 6 emerged independently from the prioritization pipeline. These candidates were not simply the most abundant or defining genes of cluster 6. For example, the TRPA1-like g25745 ranked 15th among the top 20 markers of this cluster, while many of the highest-ranked cluster 6 markers were not among the nine consensus candidates. Thus, the response to heat may involve a specific subset of the molecular program expressed by this neuronal population. Although the physiological identity of cluster 6 remains unknown, its enrichment in receptors and signal-transduction genes, together with the concentration of heat-responsive candidates, makes it a priority target for investigating how coral neurons respond to environmental change.

Importantly, several candidates converge broadly on Ca²□-dependent signaling. TRPA-family channels are nonselective cation channels that can mediate Ca²□influx, calmodulins are major intracellular Ca²□sensors, and Class A GPCRs can regulate intracellular Ca²□signaling. Although our results do not establish that these proteins operate in a common pathway, their association with the same neuronal population and coordinated downregulation under heat stress suggest remodeling of sensory and Ca²□-dependent signaling. Such downregulation need not indicate loss of sensory function. During sustained thermal exposure, reduced expression of signaling components could represent an adaptive response that alters the sensitivity or operating range of neuronal signaling, potentially limiting persistent excitation or Ca²□influx. Alternatively, it could reflect broader stress-associated suppression or disruption of neuronal function. Distinguishing among these possibilities will require direct functional measurements. Spatial localization and functional characterization, particularly of candidates concentrated in cluster 6, will be important next steps.

## Conclusion

By integrating an existing single-cell atlas with independently generated bulk RNA-seq datasets, we identified a small set of neuronal genes consistently associated with the response of *S. pistillata* to heat stress. The convergence of two complementary analytical approaches and two independent heat-stress studies reduced thousands of differentially expressed genes to nine candidates, while targeted sequence analysis refined several automated annotations and highlighted others whose molecular functions remain unresolved. These candidates provide experimentally testable entry points for investigating how coral neurons sense and respond to environmental change. More broadly, our results demonstrate that genes can be prioritized based on where they are expressed, the molecular characteristics of the cell populations in which they occur, and how consistently they respond to environmental perturbation, even when their precise molecular functions cannot be confidently inferred from homology. This framework provides a practical approach for extracting cell-type-associated biological information from the growing collection of independently generated transcriptomic resources in corals and other non-model organisms.

## Acknowledgements

We thank the Center for Advanced Research Computing (CARC) at the University of Southern California for providing the computing resources. This work was supported by the Allen Family Philanthropies.

## Data Accessibility

No new sequence data were generated in this study. Key intermediate files are available in the Figshare repository (https://figshare.com/s/fa2eda73372f74590e31). Scripts are available at GitHub (https://github.com/Ruiqi-CUB/Stylophora_Integrative_Transcriptomics).

## Author Contributions

Ruiqi Li: Conceptualization; data curation; formal analysis; investigation; methodology; visualization; writing - original draft; writing - review and editing.

Martin Tresguerres: Formal analysis; Funding acquisition; writing - review and editing.

Carly Kenkel: Conceptualization; funding acquisition; resources; supervision; writing - review and editing.

