## Supplementary figures for "Integrating single-cell and bulk transcriptomes identifies neuronal heat-stress-associated candidate genes in a reef-building coral"

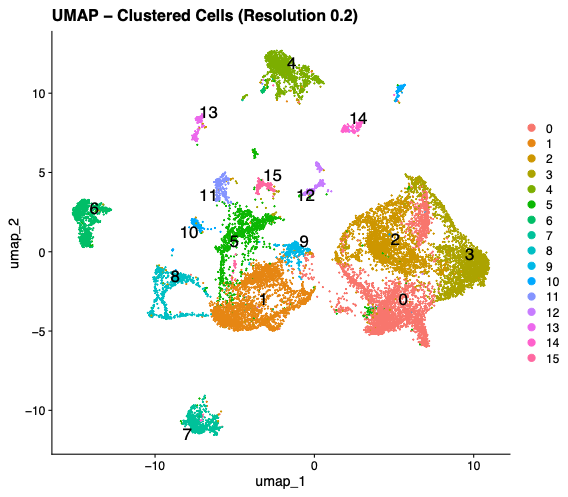

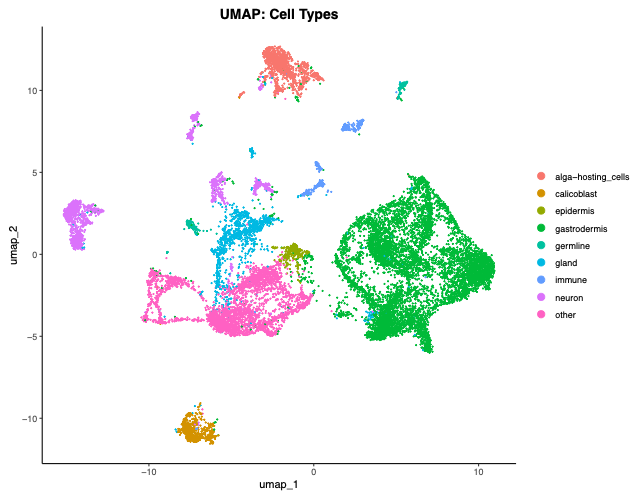

Figure S1. Single-cell clusters in *Stylophora pistillata*. UMAP visualization of single-cell transcriptomes showing the graph-based clusters (top) and their cell-type annotations (bottom). Neuronal populations were assigned to clusters 6, 11, 13, and 15 based on marker-gene expression.

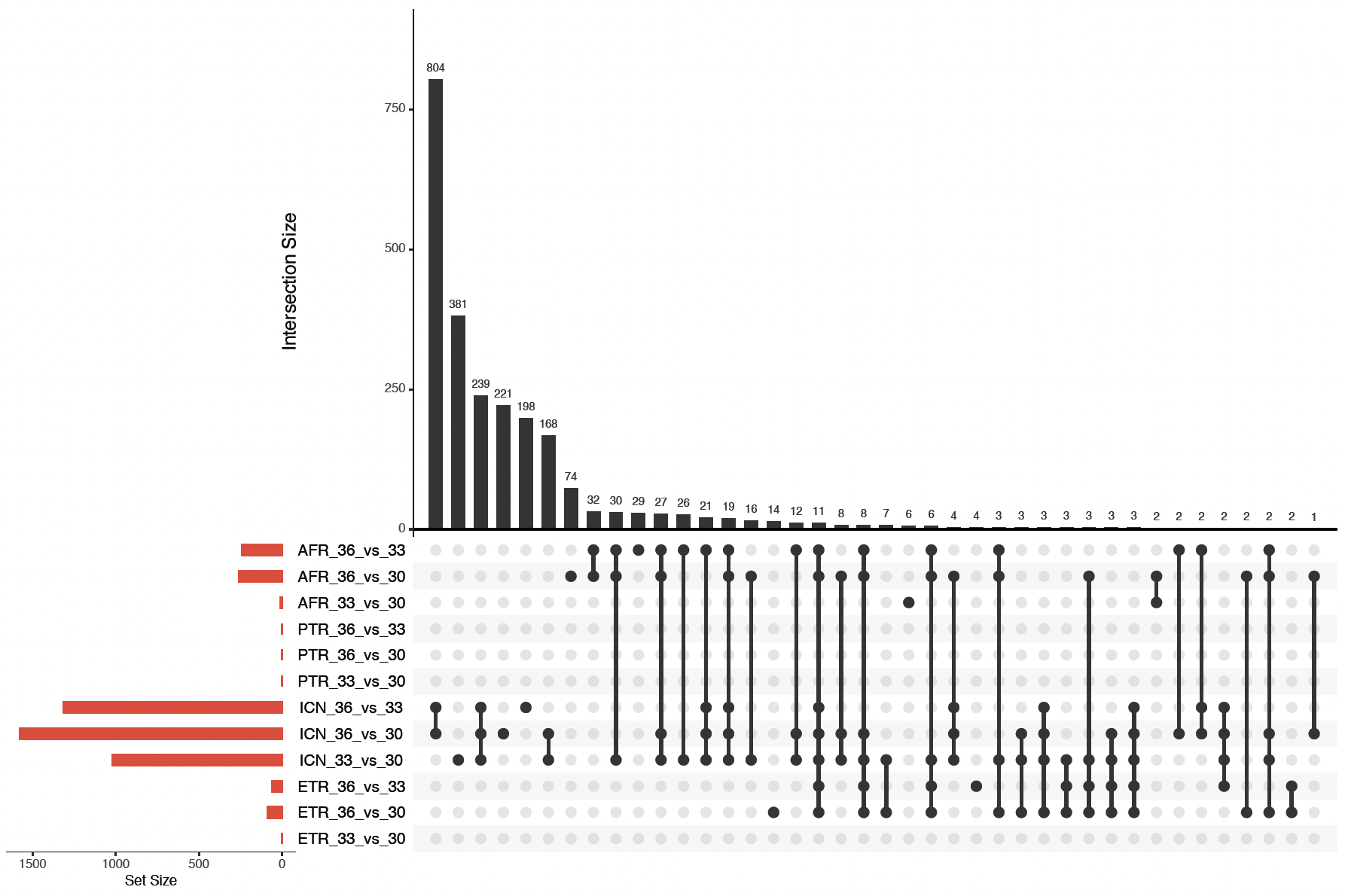

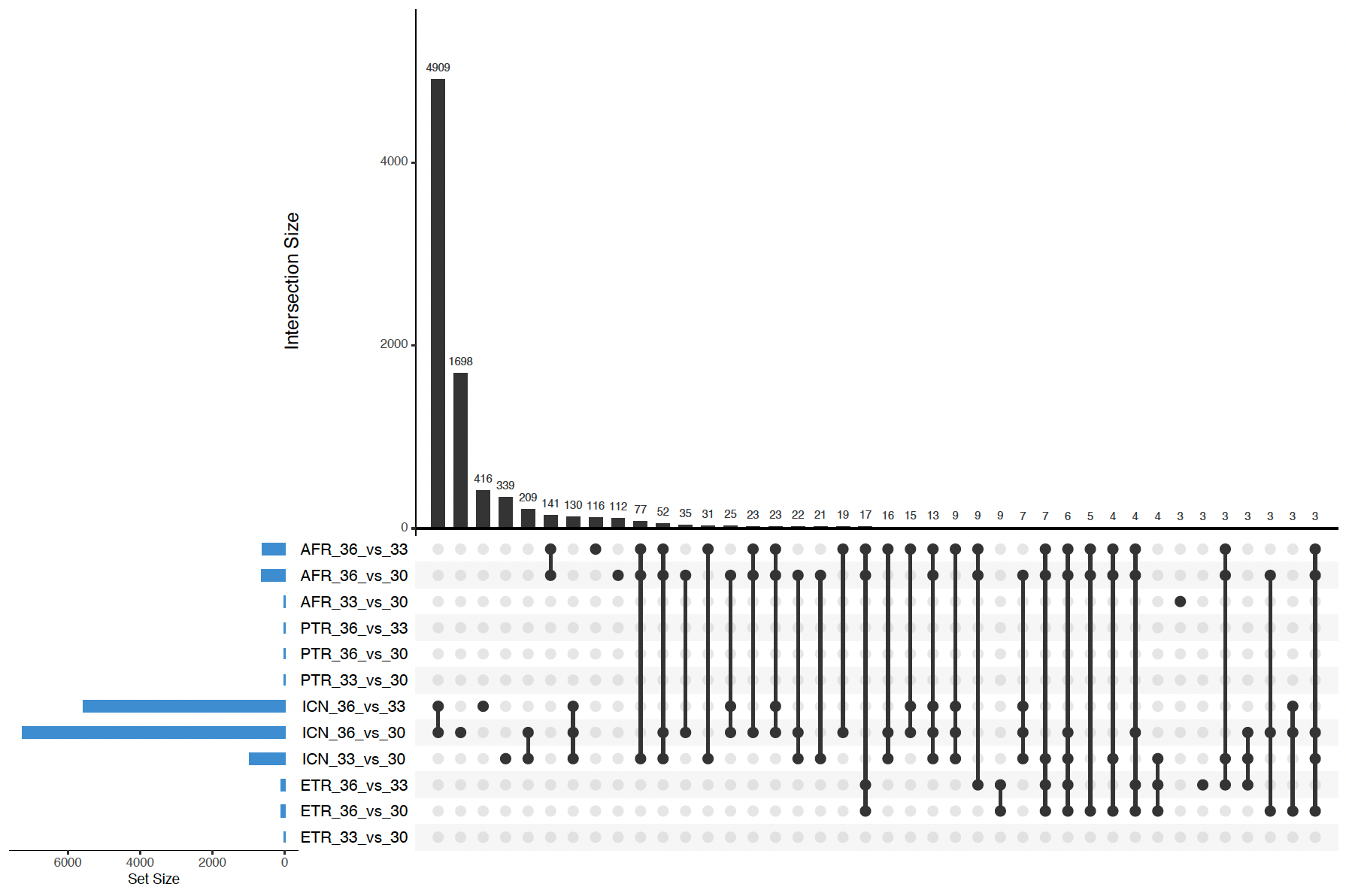

Figure S2. Overlap of differentially expressed genes (DEGs) across temperature contrasts in the Voolstra et al. (2021) dataset. UpSet plots show genes upregulated (top, red) and downregulated (bottom, blue) in comparisons among 30°C, 33°C, and 36°C within the AFR, PTR, ICN, and ETR populations. Horizontal bars indicate the size of each DEG set; connected dots identify the sets in each intersection, and vertical bars give the intersection sizes.

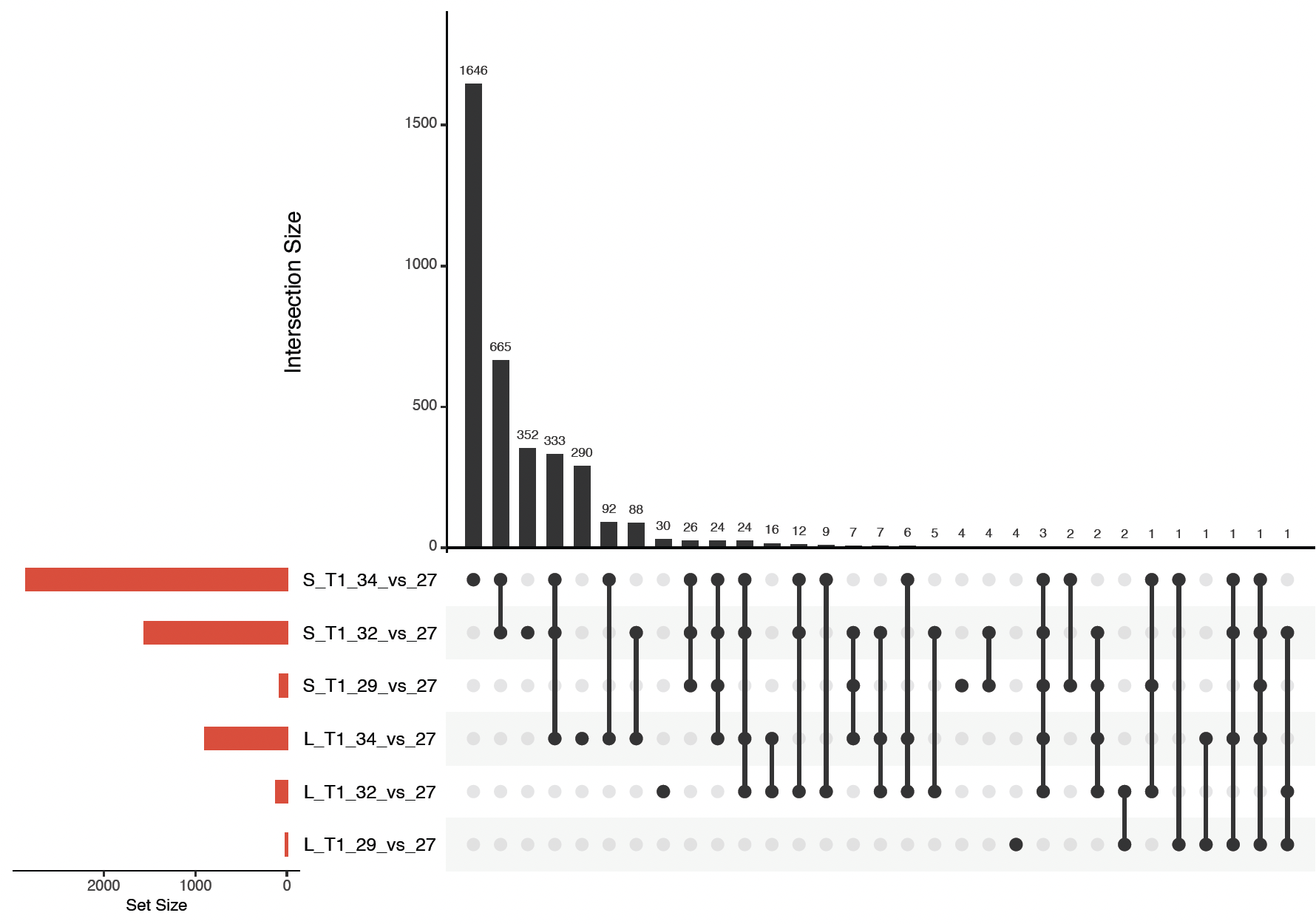

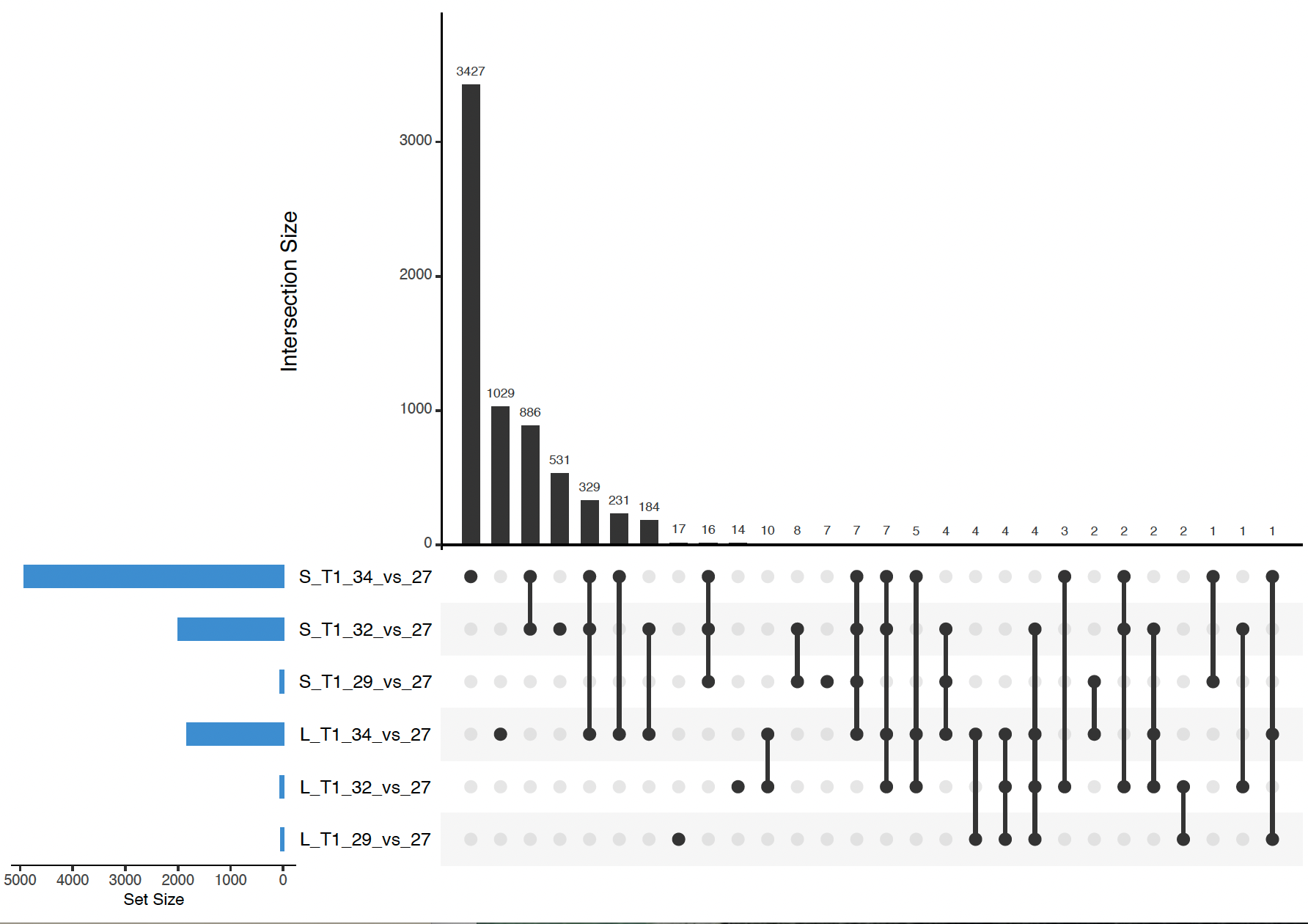

Figure S3. Overlap of differentially expressed genes across temperature contrasts in the Savary et al. (2021) dataset. UpSet plots show genes upregulated (top, red) and downregulated (bottom, blue) in the short-term (S) and long-term (L) T1 comparisons against the control treatment. Horizontal bars indicate DEG set sizes; connected dots identify the sets in each intersection, and vertical bars give the intersection sizes. Temperature contrasts are labeled in the plots.

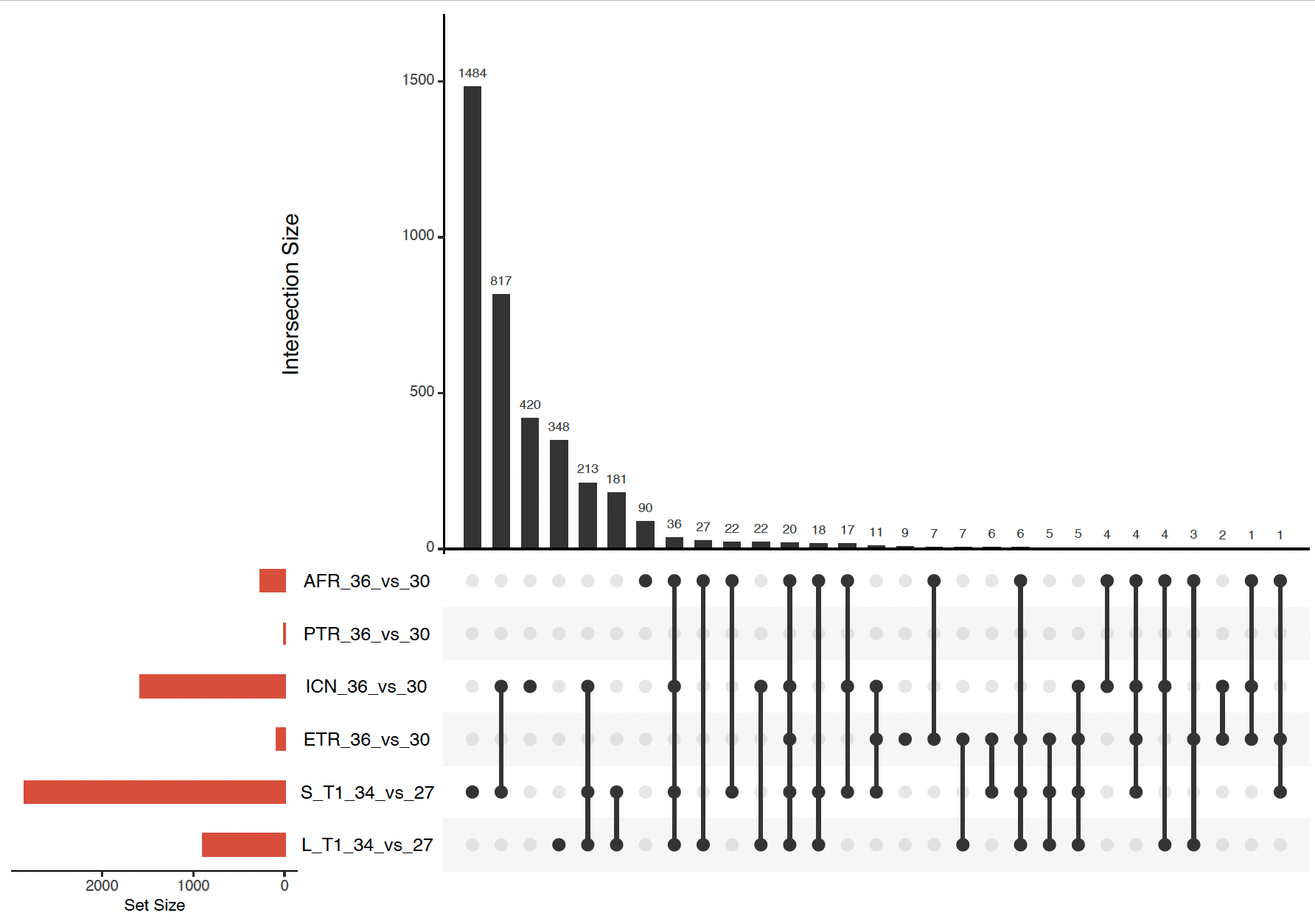

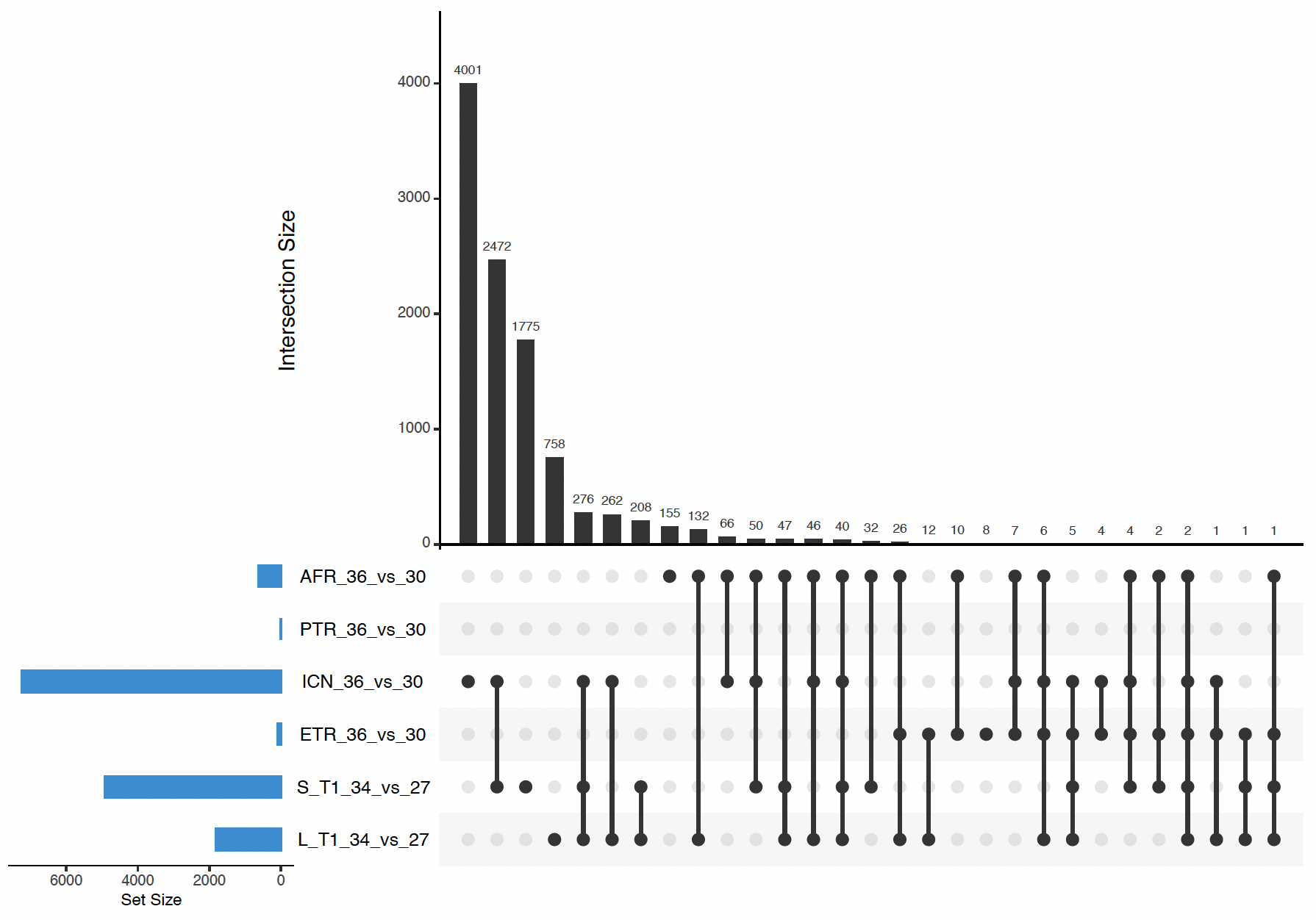

Figure S4. Cross-study overlap of heat-responsive genes. UpSet plots compare upregulated (top, red) and downregulated (bottom, blue) genes from the 36°C versus 30°C population contrasts in Voolstra et al. (2021) with the short-term and long-term T1 heat contrasts in Savary et al. (2021). Horizontal bars indicate DEG set sizes; connected dots identify the sets in each intersection, and vertical bars give the intersection sizes.

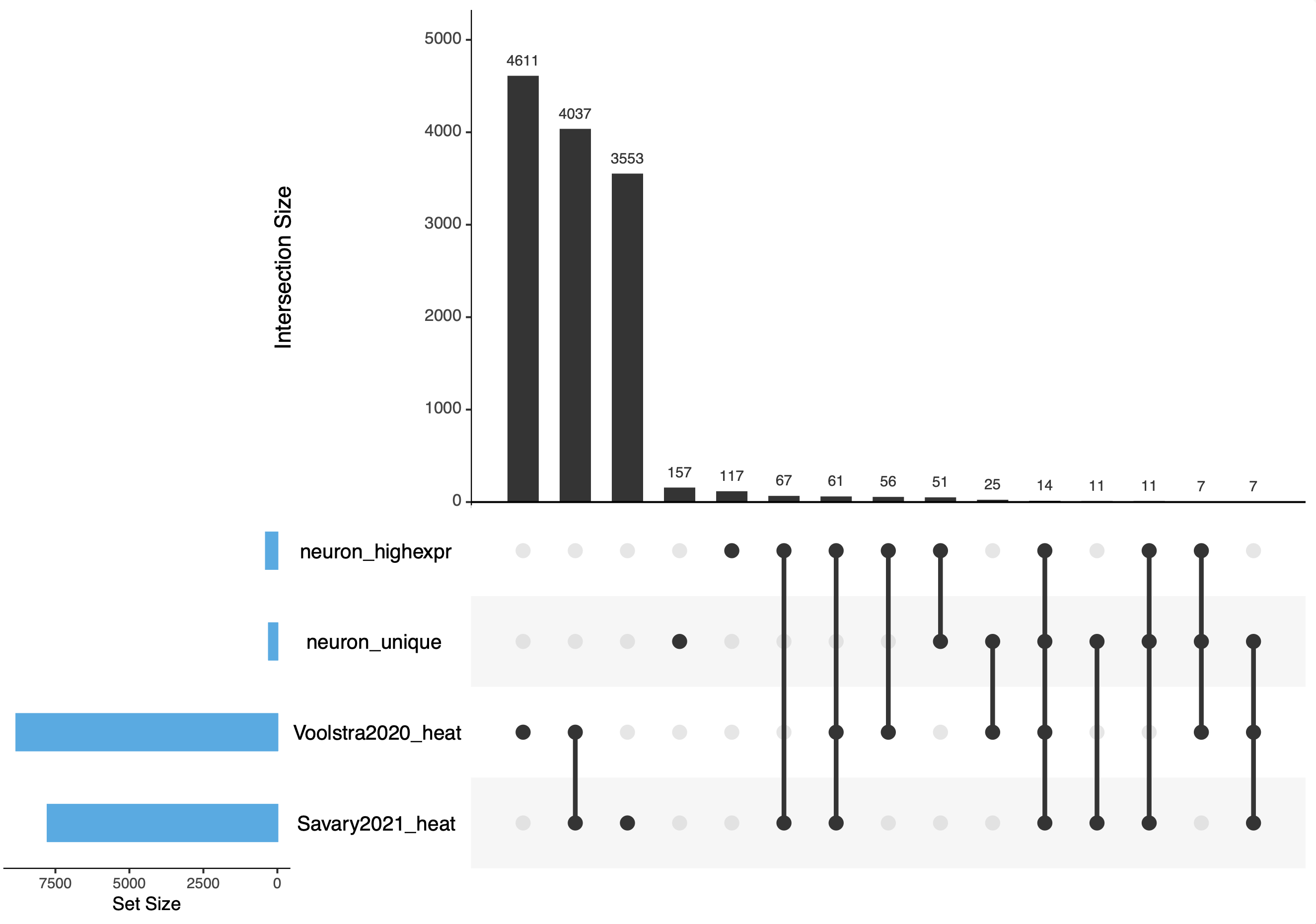

Figure S5. Intersection of neuronal gene sets with heat-responsive genes from two bulk RNA-seq studies. UpSet plot comparing genes highly expressed in neurons (neuron_highexpr), genes predominantly expressed in neurons (neuron_predominant), ICN 36°C versus 30°C DEGs from Voolstra et al. (2021), and short-term T1 34°C versus 27°C DEGs from Savary et al. (2021). Horizontal bars indicate set sizes; connected dots identify intersections, and vertical bars give the number of genes in each intersection. Fourteen genes occur in all four sets.

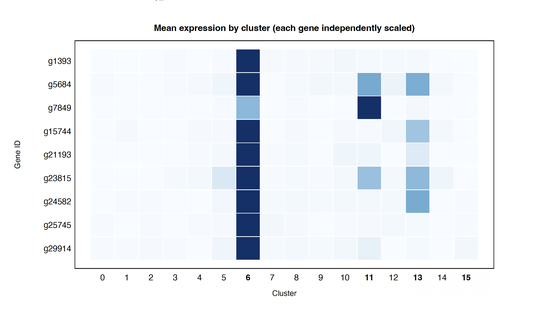

Figure. S6. Expression of nine consensus candidate genes across single-cell clusters. Heatmap of mean expression for each candidate gene across clusters in *the S. pistillata* single-cell dataset. Color intensity is scaled independently for each gene, so it shows relative expression across clusters within a row. Eight candidates have their highest mean expression in cluster 6; g7849 has its highest mean expression in cluster 11.

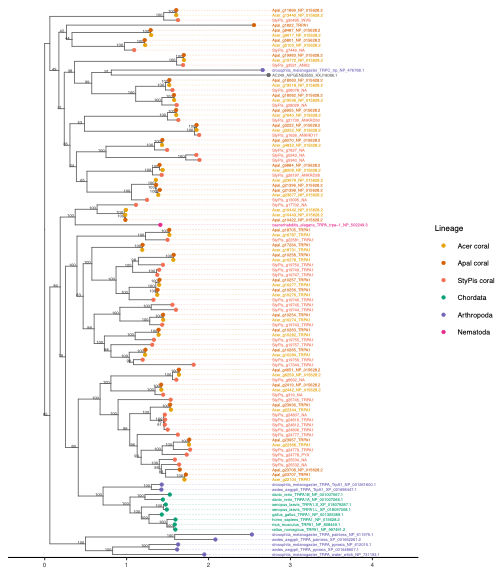

Figure. S7 Phylogenetic placement of the TRPA-family candidate g25745. Maximum-likelihood phylogeny of TRPA-family protein sequences, including g25745, proteins annotated as TRPA in the *S. pistillata* genome, additional *S. pistillata* proteome hits, and reference sequences from model species. The placement of g25745 among coral TRPA-family channels supports its characterization as TRPA-like without establishing one-to-one orthology with bilaterian TRPA1.
